# CNETML: Maximum likelihood inference of phylogeny from copy number profiles of spatio-temporal samples

**DOI:** 10.1101/2022.03.18.484889

**Authors:** Bingxin Lu, Kit Curtius, Trevor A. Graham, Ziheng Yang, Chris P. Barnes

## Abstract

Phylogenetic trees based on copy number alterations (CNAs) for multi-region samples of a single cancer patient are helpful to understand the spatio-temporal evolution of cancers, especially in tumours driven by chromosomal instability. Due to the high cost of deep sequencing data, low-coverage data are more accessible in practice, which only allow the calling of (relative) total copy numbers due to the lower resolution. However, methods to reconstruct sample phylogenies from CNAs often use allele-specific copy numbers and those using total copy number are mostly distance matrix or maximum parsimony methods which do not handle temporal data or estimate mutation rates. In this work, we developed a new maximum likelihood method based on a novel evolutionary model of CNAs, CNETML, to infer phylogenies from spatio-temporal samples taken within a single patient. CNETML is the first program to jointly infer the tree topology, node ages, and mutation rates from total copy numbers when samples were taken at different time points. Our extensive simulations suggest CNETML performed well even on relative copy numbers with subclonal whole genome doubling events and under slight violation of model assumptions. The application of CNETML to real data from Barrett’s esophagus patients also generated consistent results with previous discoveries and novel early CNAs for further investigations.

## 1 Introduction

Phylogenetic trees have been widely used in the study of cancer, providing important insights into carcinogenesis [1]. Various markers have been used for phylogeny inference, including data derived from comparative genomic hybridisation (CGH), single nucleotide polymorphism (SNP) array, fluorescence in situ hybridization (FISH), and next-generation sequencing (NGS) technologies. The rapid advances of NGS, such as whole genome sequencing (WGS) and whole exome sequencing (WES), allow the generation of huge amounts of genomic data from patient samples. NGS-derived somatic variants, mainly single nucleotide variants (SNVs) and copy number alterations (CNAs), have become common markers for phylogeny inference. CNAs are more complex than SNVs and often related to chromosomal instability (CIN) which may generate different types of structural variations (SVs) or aneuploidy [2]. Although most phylogeny inference approaches use SNVs, a number of methods are based solely on CNAs [1, 3–10]. One reason is that it is hard to detect point mutations for some cancers mainly driven by SVs or CIN [11], such as high grade serous ovarian cancer [12], oesophageal cancer [13], and osetosarcoma [14]. Another reason is that it is difficult to detect SNVs from low-coverage data whereas the larger sizes of CNAs provide more signal for reliable detection.

Given different input data and aims, trees reconstructed from CNAs called from a single patient are of four major types: 1) mutation tree when the order and evolutionary history of mutational events are of interest [15], as in SCICoNE [5] and CONET [7], where each tip represents copy number events and cells are attached to each node; 2) clone tree when clonal deconvolution is feasible, as in CNT-MD [4] and DEVOLUTION [8], where each tip represents a clone; 3) cell tree when CNAs can be called for each cell, as in FISHtree [16, 17], sitka [6], and NestedBD [10], where each tip represents a cell; 4) sample tree when each sample is assumed to be homogeneous, as in MEDICC [18], MEDICC2 [9] and PISCA [3], where each tip represents a sample.

Intra-tumour heterogeneity (ITH) causes difficulty in analysing bulk DNA sequencing (bulk-seq) data, where only the aggregated signals can be observed. Therefore, phylogeny inference from bulk-seq data is often coupled with clonal deconvolution that determines the number and fraction of clones in a sample [1]. Reliable quantification of subclonal CNAs and ITH requires deep sequencing on samples of good quality and is expensive. Single cell DNA sequencing (sc-seq) circumvents the need to infer clone structure, but the data are still very noisy [15, 19]. Low-coverage bulk-seq, such as shallow WGS (sWGS), are instead more cost-effective and accessible, especially for SV-driven cancers [20]. They have been widely applied to detect CNAs, particularly on formalin-fixed paraffin-embedded (FFPE) samples, which are commonly available for diagnostics but have low DNA quality [11, 13, 21–23], and cell-free DNA in plasma [24, 25]. There have been sWGS data for patient samples taken over time and space during a long time period, such as in the surveillance of Barrett’s esophagus (BE) [13] and inflammatory bowel disease (IBD) [23], and potentially more longitudinal samples will be sequenced from both pre-cancers and tumours of the same individual [26]. These longitudinal samples also contain temporal information that can be used to estimate node ages and mutation rates, which are important parameters in carcinogenesis. However, only a few reliable methods exist to detect CNAs from sWGS data, especially absolute copy numbers [27]. Most of the previous sample phylogeny inference methods are designed for absolute allele-specific integer copy numbers which are often called from SNP arrays and high-coverage NGS data, such as MEDICC [18], MEDICC2 [9], and PISCA [3]. To better understand cancer progression from these sWGS data, it is important to have methods that can build sample trees based solely on (relative) total copy numbers, which will be addressed in this paper.

The model of CNA evolution is critical for phylogeny inference, but it is challenging to propose a model which maintains a good trade-off between biological realism and complexity [19]. The underlying mechanisms of CNAs are often very complicated, such as chromothripsis, breakage fusion bridges, and failure in cell cycle control [22]. As a result, CNAs vary from small focal duplication/deletion to chromosome-level gain/loss and whole genome doubling (WGD) at different rates [28], which creates complex dependencies across the genome, such as overlaps, back mutations, convergent and parallel evolution [29]. Therefore, the infinite sites or perfect phylogeny assumption, which is commonly used in inferring phylogeny from SNVs, is often violated, as is the infinite alleles or multi-state perfect phylogeny assumption [19]. The models for genome rearrangement, microsatellite, and multigene families seem relevant yet hard to transfer to CNAs [19, 30].

Some methods transform original copy number calls into presence or absence of changes (break-points) [6, 31], which are less likely to overlap, so that the infinite sites assumption is well approximated. Although this representation simplifies the complex spatial correlations across sites, it does not use the full copy number data. Other methods represent the genome as a vector of copy number values, often called copy number profile (CNP) [4]. Based on CNPs, some methods build trees without a model, such as the maximum parsimony (MP) method with the Fitch algorithm [23, 32] and distance matrix methods based on Euclidean [33] or Manhattan [34] distances, and hence they may underestimate the true evolutionary distance as no correction of hidden changes is applied [9, 31]. Other methods use copy number transformation (CNT) models that allow the computation of minimum evolutionary distance between CNPs, which is the shortest sequence of events that transform one CNP to the other. One such model was implemented in FISHtree [16, 17], which assumes each event (single gene gain/loss, chromosome gain/loss, or WGD) affects a single unit (gene or chromosome or genome) independently, with or without weights for different types of events. Another well-studied model, within MEDICC [18], assumes an event (segment duplication/deletion) may affect contiguous segments of variable size. This model deals with horizontal dependencies caused by overlapping CNAs and hence is less likely affected by convergent evolution. It has been extended to allow weights on CNAs of different position, size, and type (duplication/deletion) [35] and WGD [9, 36]. The weighted versions of both models allow the estimation of CNA rates in term of event probabilities [17, 35], but mutation rates by calendar time cannot be estimated. A few CNP-based methods use the finite sites models, or continuous-time Markov chains, which have good theoretical properties and are frequently used to model nucleotide changes [37]. Although Markov chains often assume independent sites to simplify computation, which is violated by overlapping CNAs, it corrects multiple hits at the same site and serves as a workable model of CNAs. For example, SCONCE used a Markovian approximation that combines the temporal Markov process with a spatial hidden Markov model (HMM) to detect CNAs in sc-seq data [38]; Elizalde et al. used the product of 23 Markov chains to model numerical CIN of individual chromosomes in clonally expanding populations [39]. Markov model makes it possible to use statistical methods to infer CNA-based trees, mutation rates by time, and ancestral genomes, such as maximum likelihood (ML) method and Bayesian method. PISCA used such a Markov chain to model gain, loss, and conversion of haplotype-specific copy numbers called from SNP array data [3]. NestedBD used a birth-death model, a special type of Markov chain where transitions from state *i* can only go to state *i* + 1 or *i* − 1, for total copy numbers called from sc-seq data, where a birth (death) event corresponds to copy number amplification (deletion) [10]. Both PISCA and NestedBD are implemented as packages in the popular Bayesian evolutionary analysis platform BEAST [40, 41], and hence are not easily adapted for more bespoke mutation models that will be required for understanding carcinogenesis. In addition, most phylogeny inference methods based on CNPs cannot handle multiple scales of chromosomal changes due to the inherent complexity. Notable exceptions are FISHtree, which was designed for FISH data and is not scalable for longer CNPs [16, 17], and MEDICC2 which only considered segment duplication/deletion and WGD but excluded chromosome or arm level gain/loss [9].

In this paper, we developed an approach based on a novel Markov model of duplication and deletion, CNETML, to do maximum likelihood inference of single patient phylogeny from total copy numbers of spatio-temporal samples. To the best of our knowledge, this is the first method to jointly infer the tree topology, node ages, and mutation rates of temporal patient samples from (relative) total CNPs called from sWGS data. CNETML is applicable to haplotype-specific CNPs as well, which is the basis of our model and considered as missing information when total CNPs are taken as input. We also developed a program to simulate CNAs from patient samples, CNETS (Copy Number Evolutionary Tree Simulation), which was used to validate sample phylogeny inference methods. Our programs are mainly implemented in C++, available at https://github.com/ucl-cssb/cneta. The results on extensive simulations suggest that CNETML accurately recovered the tree topology, node ages, mutation rates, and ancestral CNPs when there were sufficient CNAs and sampling time differences present in the data. CNETML on total CNPs performed as well as haplotype-specific CNPs when less than 10% of copy-neutral CNAs existed in the simulated data. CNETML also had good accuracy when applied to relative CNPs from simulated data with subclonal WGDs, which is desirable for applications to sWGS data. Moreover, the simulations suggest CNETML was robust to slight violations of model assumptions and that it obtained reasonable inferences on data of typical focal CNA size. We applied CNETML on relative CNPs called from two BE patients in existing literature and obtained results consistent with previous findings and novel early CNAs from reconstructed ancestral CNPs which are worth further validations, suggesting the utility of CNETML.

## 2 Results

### Overview of CNETML

The input of CNETML (Figure 1) includes a set of integer total/haplotype-specific CNPs for multiple samples of a patient and/or sampling timing information (in year) if available (see section 4 for details on input preparation). The length of each CNP is the number of sites in a sampled genome, which can be either bins or segments, and we assume all genomes have the same sites. Here, a bin is a genomic region of fixed size and a segment is a genomic region of variable size which may be obtained by merging consecutive bins with the same copy number. In CNA detection, the general steps include binning, bias removal, segmentation, and copy number assignment [19, 42]. In binning, the genome is divided into bins of certain size, usually fixed, and reads aligned to each bin are counted. In segmentation, the genome is partitioned into a series of segments whose copy number is different from that of the adjacent segment. Therefore, although a site is not as well defined as when modelling SNVs on individual nucleotides, it is feasible to consider a bin or a segment as a site.

**Figure 1:**
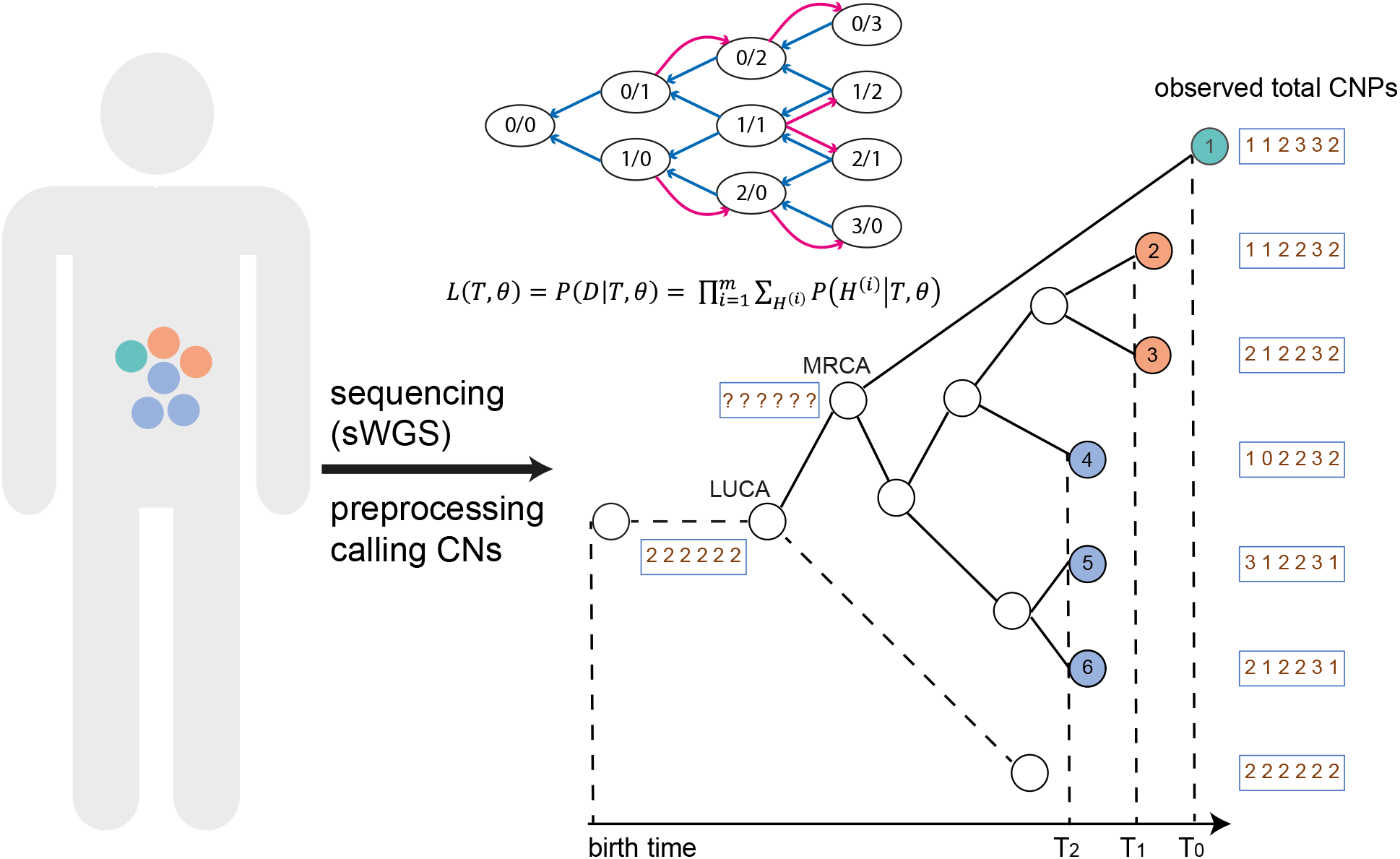
The schematic overview of CNETML. Samples may be taken from a patient at different locations and times during surveillance and get sequenced (with sWGS) and analyzed to generate copy numbers (CNs). Given the CNPs and/or sampling times of all the patient samples, CNETML aims to infer a sample tree in which tips correspond to observed CNPs in samples and internal nodes correspond to ancestral CNPs. From the root, which represents the last unaltered common ancestor with normal copy number state (LUCA), there is a branch of length zero (dashed line), which leads to a tip representing the normal CNP to get a binary tree. LUCA is connected to the most recent common ancestor (MRCA) of the patient samples. We added an additional node before LUCA to show the CNP at the birth time, which was used to constrain the age of LUCA in inference. The state transition diagram of the Markov chain shows the duplication (red arrow) and deletion (blue arrow) of haplotype-specific copy number, with the maximum total copy number being 3 and the value in each oval representing a possible combination of copy number for haplotype A and B, denoted by *c_A_/c_B_*. The Markov model allows the computation of tree likelihood by taking the product over all sites along the genome. The CNPs of internal nodes (including MRCA) are unknown and inferred with ancestral reconstruction algorithms. Samples taken at different time points (*T*_0_, *T*_1_, and *T*_2_) are denoted by different colours.

We treat an integer copy number at each site as a discrete trait whose states are dependent on the maximum possible copy number. To maintain model simplicity, we assume copy numbers at the sites of a genome change independently of each other (independent sites assumption) and the change of copy number at each site follows a continuous-time non-reversible Markov chain. The Markov chain naturally starts from the normal diploid copy number and has an absorbing state when no copy remains. Due to the difficulty in incorporating CNAs of different scales, we propose a model of site duplication and deletion at haplotype-specific level, which is similar to that in PISCA [3] yet designed for processing total CNPs. Moreover, we consider CNA rate (or mutation rate) per haplotype per site per year and allow user-specified maximum copy number.

Suppose *c_max_* is the maximum total copy number, then each site has *S* possible states {0, 1, 2, …, *S*−1}, where

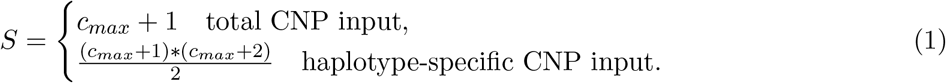

The change of haplotype-specific copy numbers on each site via duplication (deletion) at rate *u* (*e*) per haplotype per site per year is specified by the rate matrix *Q* (see Supplementary Table 1 for *Q* at *c_max_* = 4). In *Q*, we list haplotype-specific copy numbers in order of increasing total and haplotype *A* copy number so that each combination of *c_A_* and *c_B_*, (*c_A_, c_B_*), corresponds to a unique state, where *c_A_* and *c_B_* represent the copy numbers for two haplotypes respectively. For example, normal copy number (1, 1) is represented by state 4, and copy number (4, 0) is represented by state 14. Note that (*c_A_, c_B_*) and (*c_B_, c_A_*) are distinguishable in the data when haplotype-specific copy numbers are provided. Suppose a genome *j* has *m* sites and *c_ij_* is the copy number state at site *i*, which is either the total copy number or the state corresponding to the haplotype-specific copy number in *Q*. Then its observed CNP is denoted by (*c*_1*j*_, *c*_2*j*_, …, *c*_*mj*_). The CNPs for all the *n* sampled genomes form a data matrix of *n* rows and *m* columns, denoted by *D*. The observed copy number states across all samples at a site *i* is called a site pattern, denoted by 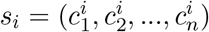. We say site *i* is invariant if *s_i_* is composed of normal copy number states only, and variant otherwise.

The likelihood for a tree *T* of *n* samples with parameters *θ*, *L*(*T, θ*), is the probability of observing *D* at the tips of *T* given *θ*. The Markov model specified by *Q* allows the computation of *L*(*T, θ*) by taking the products of probabilities at individual sites:

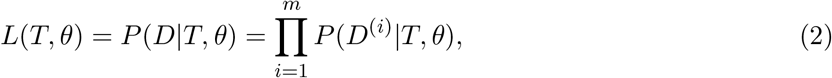

where *D*^(*i*)^ is the *i_th_* column of *D*. When taking total CNPs as input, we revise *L*(*T, θ*) to incorporate haplotype-specific copy numbers as missing information, which is similar to the handling of ambiguities in a nucleotide substitution model [37]:

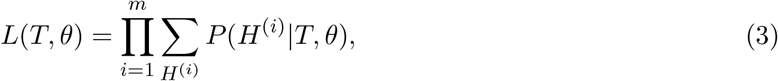

where *H* is a data matrix of unknown haplotype-specific copy number states that are compatible with *D*, *H*^(*i*)^ is the *i_th_* column of *H*, and there may be multiple such matrices for *D*. For example, the probability of observing total copy number 3 is a sum over all compatible haplotype-specific copy numbers (0, 3), (1, 2), (2, 1), and (3, 0). In other words, the haplotype-specific copy numbers are latent variables, and the likelihood is an average over them.

We computed *L*(*T, θ*) with Felsenstein’s pruning algorithm [43] with a few adaptations described in section 4. *L*(*T, θ*) was maximised by minimizing its negative logarithm function with L-BFGS-B algorithm [44], a numerical iterative method with bound constraints. Due to the super-exponentially increasing number of trees with the number of tips, we implemented two approaches to search the tree space and get the ML tree. One is exhaustive search which enumerates all the possible tree topologies for trees of less than seven samples. The other is heuristic search for larger trees, adapted from the approach in IQ-TREE [45], a popular ML phylogeny inference program.

When there are samples taken at different times, it is feasible to estimate mutation rates according to the differences of CNPs and sampling times, similar to the dating of virus divergences [37]. Although mutation rates during neoplastic progression are likely to change over time due to CIN [3], it is unlikely that they can be estimated reliably, so we assumed constant mutation rates under a global clock for simplicity. We jointly estimated the tree topology, mutation rates, and node ages (starting from 0 at birth time) with the following constraints in optimization: 1) The age of each internal node must be smaller than all its descents; 2) The age of root node must be smaller than the patient age at the first sample time or the tree height in year is smaller than the patient age at the last sample time. We transformed node age variables to encode the constraints imposed by patient ages at different sampling times so that *θ* = (*x*_1_, *x*_2_, …, *x_n_, u, e*), where *x_i_* is the transformed variable for age of an internal node *i* and converted back to branch length in year later (see section 4 for more details). When all the samples are taken at the same time, there is no information to estimate mutation rates and node ages at the same time, so *θ* = (*l*_1_, *l*_2_, …, *l*_2*n*−1_), where *l_i_* = (*u*_0_ + *e*_0_)*t_i_* is the length of branch *i* measured by expected number of CNAs per site, *u*_0_ (*e*_0_) is the pre-specified duplication (deletion) rate per haplotype per site per year, and *t_i_* is the time in year covered by branch *i*. Here, we separate (*u*_0_ + *e*_0_) and *t_i_* for the convenience of implementation within CNETML.

Ancestral reconstruction may suggest early CNAs that are likely cancer driver events and useful for early diagnostics. Therefore, we reconstructed ancestral states at variant sites with unique site patterns based on the obtained ML tree using classical methods, including both marginal reconstruction of the most recent common ancestor (MRCA) node and joint reconstruction of all ancestral nodes [46, 47]. For marginal reconstruction, we computed the posterior probability of each possible copy number state for MRCA and assigned the state with highest probability to each site. For joint reconstruction, we assumed the best reconstruction is obtained when the root has normal diploid copy number states.

We used bootstrapping to measure the uncertainties of an estimated ML tree *T_m_* [37]. To get a bootstrap tree, we sampled sites from the input data matrix *D* with replacement to get a pseudo-sample *D*′ with the same dimension as *D* and built a ML tree from *D*′. The branch support value in *T_m_* is defined as the percentage of bootstrap trees including this branch (split) and computed with function pro.clade in R library ape [48].

### Validation on simulated data

#### Data simulation and comparison metrics

To validate CNETML, we developed CNETS to simulate CNAs along a phylogenetic tree of multiple patient samples (Figure 2, see section 4 for more details). In CNETS, we first generated a coalescence tree to represent the genealogical relationships among samples, the subtree starting from MRCA, under either the basic coalescent or an exponential growth model with rate *β*. We then added another node before MRCA to represent the last unaltered common ancestor with normal copy number state (LUCA) and a branch of length zero from LUCA to a new tip which represents a normal genome to obtain a binary tree. The time from LUCA to MRCA was sampled from an exponential distribution with rate which was either based on the exponential growth rate *β* or sampled from an uniform distribution 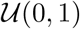. To get different sampling times, we increased the terminal branch lengths by random integer multiples of *dt* (in year), with the maximum multiple being the number of samples. We implemented two modes of simulating CNPs which differ in the types of CNAs and recorded details. When only site-level CNAs are considered and the exact mutational events are not of interest, CNPs were simulated directly along each branch of the tree according to the rate matrix *Q* with each site being a segment of variable size [37]. When CNAs of multiple scales are considered, events were simulated by generating exponential waiting times with each site being a bin of fixed size (500 Kbp by default), which allows more complex models of evolution and the recording of more detailed information for each event. CNETS generates files that record haplotype-specific/total CNPs, sampling times in year, tree topology, and CNAs along the branches respectively. The simulated CNPs at the tips and/or the tip timing information serve as input for CNETML.

**Figure 2:**
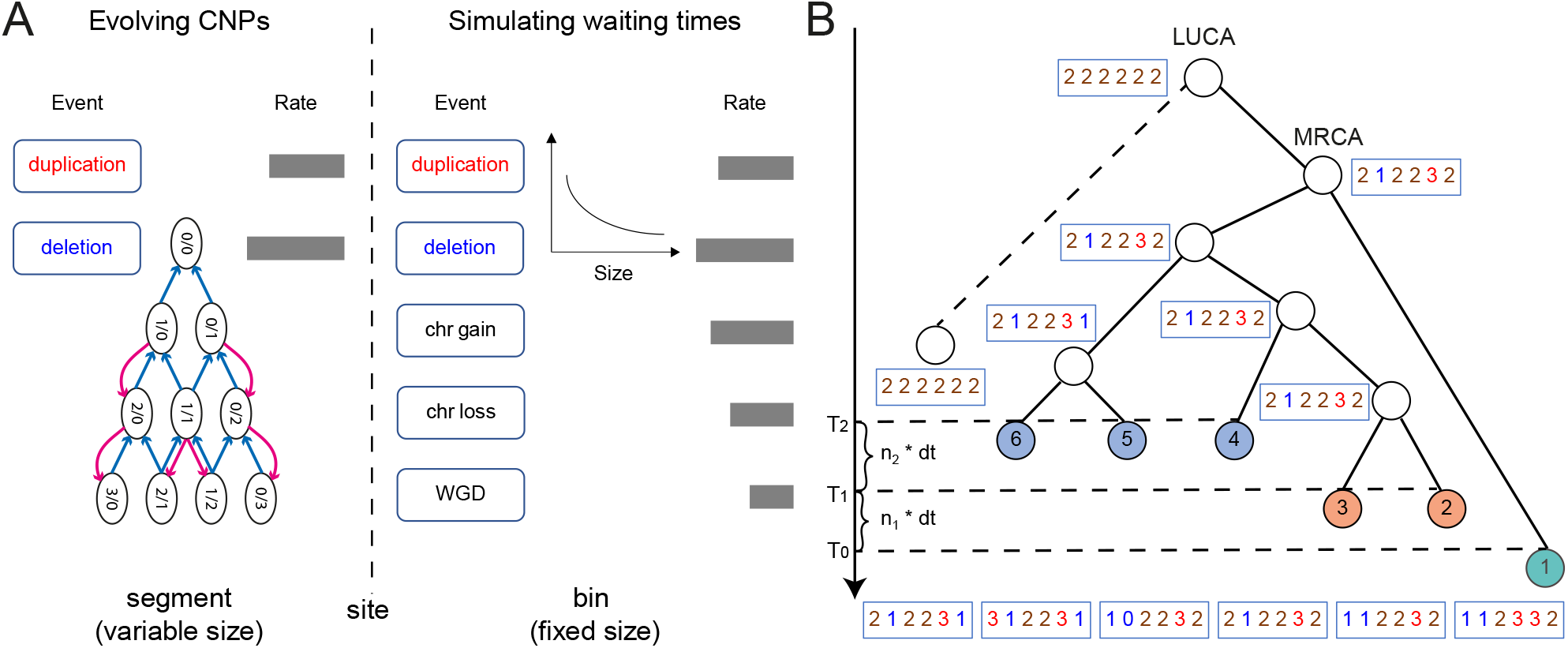
The schematic overview of CNETS. **A**: Two modes of simulation implemented in CNETS. One is simulating CNPs directly for site duplication/deletion based on segments of variable size, which follows exactly the Markov model used for phylogeny inference. The other is simulating waiting times for events of multiple scales based on bins of fixed size, in which at most five types of events (site duplication/deletion, chromosome gain/loss, and WGD) are allowed and the duplication/deletion size in the number of bins is sampled from an exponential distribution of the user-specified mean size. **B**: The simulated tree and CNPs (red: duplication, blue: deletion, brown: normal), where coloured tips represent patient samples taken at different time points *T*_0_, *T*_1_ = *T*_0_ + *n*_1_ ∗ *dt*, and *T*_2_ = *T*_1_ + *n*_2_ ∗ *dt*, with *dt*, *n*_1_, and *n*_2_ being integers and 1 <= *n*_1_, *n*_2_ <= 6 (the number of samples).

In tests, we simulated trees with parameters used in [3], which approximate an exponentially expanding haploid cancer cell population with MRCA being 20 years from the present (Supplementary Table 2). To ensure that the model used for simulation and phylogeny inference are the same, we used the simulation mode of evolving CNPs when only site-level mutations were considered. We simulated trees with *n* = 5 samples when not testing the performance of tree searching, as it is fast to enumerate all the possible trees for such small trees. Without loss of generality, we set *c_max_* = 6 and used the same rates for duplication and deletion. To get a reasonable range of mutation rates suitable for phylogeny inference, we performed tests with *u* = *e* ∈ {0.0001, 0.001, 0.01, 0.1, 1} (per haplotype per site per year) (Supplementary Figure 1). This analysis suggests that intermediate rates {0.001, 0.01} (per haplotype per site per year) are more informative for phylogeny inference, which were used in our subsequent tests.

To quantify the differences of both topologies and branch lengths between the simulated (true) tree and the inferred tree, we computed normalized Robinson–Foulds (RF) distance [49] and branch score distance [50], with smaller values indicating more accurate estimation. These distances were computed with function treedist in R library phangorn [51]. The branch length was measured by time in year in the computation of branch score distance. When the mutation rates were not estimated, *u*_0_ and *e*_0_ were set to be real values used for simulation. We also computed the differences between the estimated and true values of duplication/deletion rates and LUCA age to check the accuracy of their estimation. To measure the accuracy of ancestral reconstruction, we computed the fraction of correctly recovered states over the number of variant sites with unique site patterns for each internal node and the mean fraction over all internal nodes under joint reconstruction.

#### Performance on reconstructing trees and ancestral states

In principle, ML phylogeny inference is consistent, which means that the ML tree will converge to the true tree when the size of the data (the number of sites) increases [37]. To check the consistency of CNETML, we applied it on data simulated with different number of sites and mutation rates. To reduce confounding effects, we simulated trees with *n* = 5 samples at the same time and did not infer mutation rates. As shown in Figure 3A, all the simulated trees were better recovered with more sites and higher mutation rates, which confirms the consistency of CNETML. In the subsequent simulations, we fixed the number of sites *m* = 1000 when not stated.

**Figure 3:**
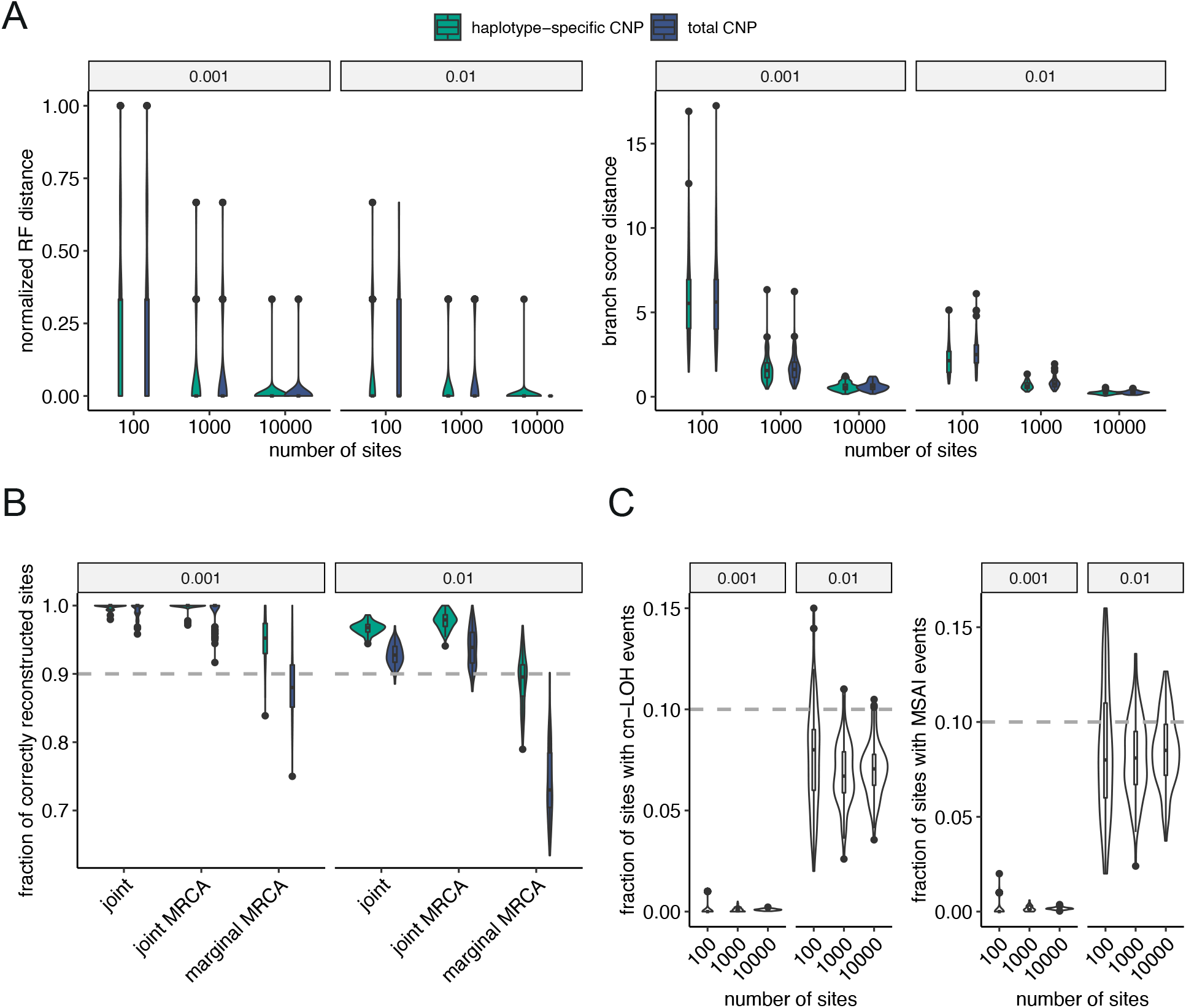
The performance of CNETML on reconstructing trees and ancestral states with total or haplotype-specific CNPs when samples are taken at the same time. **A**: The accuracy of phylogeny inference on data simulated with different number of sites and mutation rates. **B**: The accuracy of ancestral state reconstruction on simulated data with 1000 sites under different mutation rates. **C**: The fraction of cn-LOH and MSAI events in the simulated data. There are five samples in each simulated tree and 100 datasets for each parameter setting. The plots are grouped by mutation rates. The box plots show the median (centre), 1st (lower hinge), and 3rd (upper hinge) quartiles of the data; the whiskers extend to 1.5× of the interquartile range (distance between the 1st and 3rd quartiles); data beyond the interquartile range are plotted individually.

We showed that the heuristic tree search algorithm performed well on simulated data with 10, 20, and 30 samples under different mutation rates (Supplementary Figure 2). Given more samples, the estimated tree topologies did not deteriorate much, whereas the branch score distances to the true trees slightly increased. As expected, the reconstructed trees were more similar to the ground truth with more mutations.

We also checked the performance of CNETML on reconstructing ancestral states on the simulated data with 1000 sites under different mutation rates by supplying the simulated true tree and real mutation rates as input. The results suggest that more than 90% of the unique variant sites were accurately reconstructed, except when doing marginal reconstruction (Figure 3B). The fraction of accurately reconstructed sites decreased with larger mutation rate due to the presence of more variant unique sites. Joint reconstruction appeared more accurate, probably because it computes the joint probability of all the internal nodes [37].

With total copy numbers, copy-neutral loss of heterogeneity (cn-LOH) and mirrored subclonal allelic imbalance (MSAI) events (CNAs affecting different alleles of the same sites in different samples) cannot be detected. To see how total CNPs impact the inference, we applied CNETML on haplotype-specific CNPs and found that the results were not largely different except when there were more than 10% sites with cn-LOH or MSAI events (Figure 3A,C). However, the accuracy of reconstructing ancestral states was better with haplotype-specific CNPs (Figure 3B). The analysis on PCAWG dataset [52]	shows that around 80% samples have no more than 10% of the genome with cn-LOH (Supplementary Figure 3), and hence these results suggest that total CNPs can provide good approximations in practice despite information loss.

#### Performance on jointly estimating the tree and mutation rates

One major utility of CNETML is to jointly estimate the tree topology, node ages, and mutation rates when the samples were taken at different time points. The reliability of rate estimation depends on the extent of time differences at the tips, with larger differences providing more information for inference [53]. Since the sampling time differences for a patient may range from one year to 15 years as in a BE dataset [13], we simulated data under different temporal signal strengths, *dt* ∈ {1, 3, 5} (years), where the range of simulated sampling times approximated real data and samples simulated under a larger *dt* generally had larger time differences (Supplementary Figure 4). We also simulated data with 10,000 sites to check the consistency of CNETML during joint estimation. We grouped the simulated data by the mean pairwise absolute difference of tip relative times (denoted by *T_m_*) into three groups: “small” difference when *T_m_* < 3, “intermediate” difference when 3 <= *T_m_* < 7, and “large” difference when *T_m_* >= 7. The numbers of samples in each group are shown in Supplementary Table 3. Because the L-BFGS-B optimization algorithm is iterative, the initial values of parameters 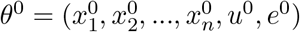 are required, where 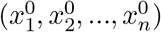 is derived from the initial tree (see section 4 on how to get initial trees)) and (*u*^0^, *e*^0^) has to be specified manually. Since the L-BFGS-B algorithm may converge to a local peak on the likelihood surface of a tree, we tried different initial values, *u*^0^ = *e*^0^ ∈ {0.0005, 0.001, 0.005, 0.01} (per haplotype per site per year), and found that CNETML was robust, except when the real mutation rate was low (0.001) and a high initial mutation rate (0.005 or 0.01) was supplied (Supplementary Figure 5). Therefore, we recommend starting from smaller initial mutation rates in real data when the range of rates is unknown and reported the results with *u*^0^ = *e*^0^ = 0.0005 (per haplotype per site per year) in Figure 4.

**Figure 4:**
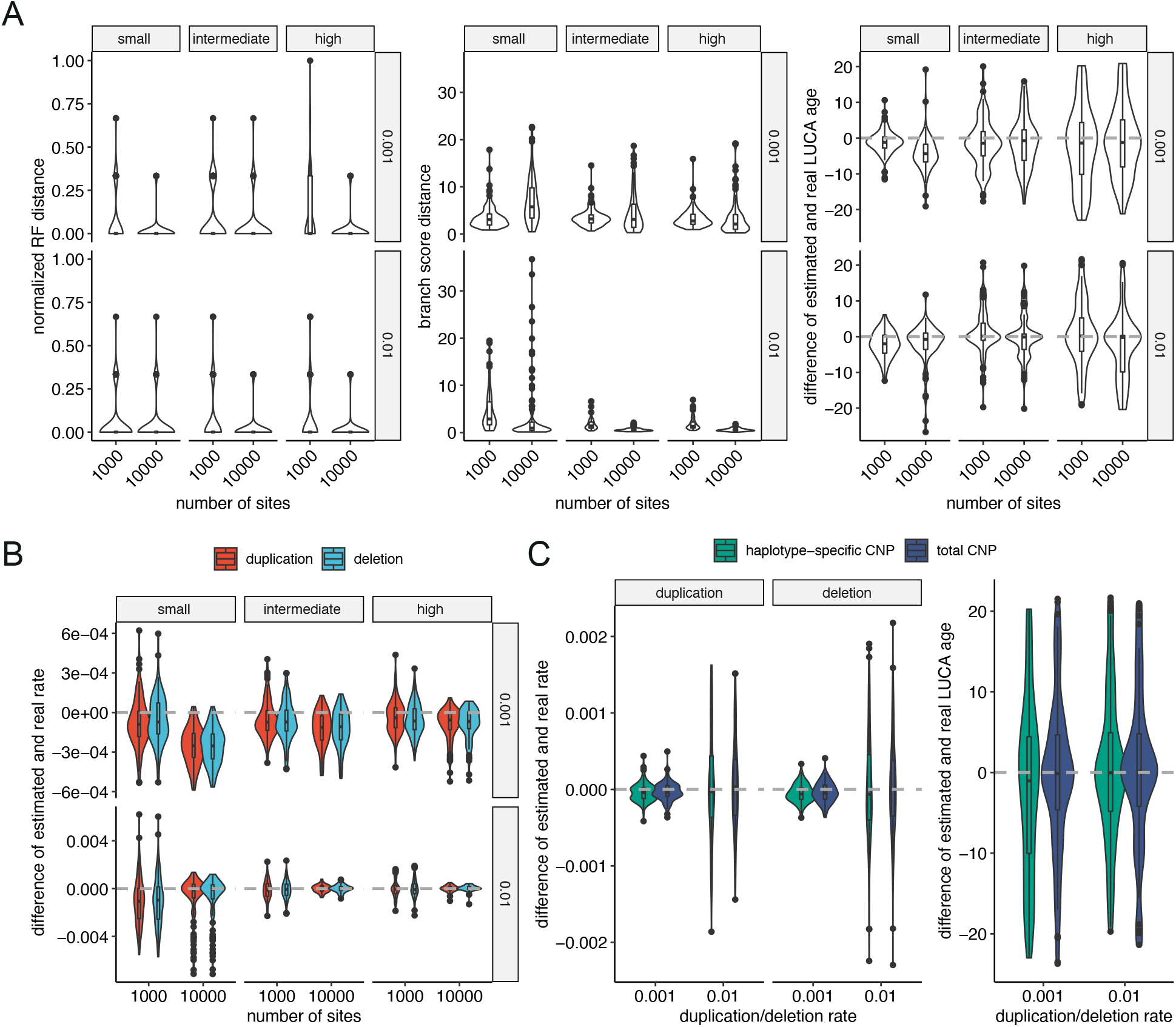
The performance of CNETML on jointly estimating the tree topology, node ages, and mutation rates on data simulated with different sampling times and mutation rates. **A**: The accuracy of phylogeny inference. **B**: The accuracy of mutation rate estimation. **C**: The accuracy in estimation of the mutation rate and LUCA age with total or haplotype-specific CNPs on simulated data with *dt* = 5 years. There are five samples in each simulated tree and 100 datasets for each parameter setting. Grey dashed line: real values. Box plots as those in Figure 3.

The joint estimation was generally better with higher mutation rates and a larger number of sites (Figure 4A-B). As shown in Figure 4A, the sampling time differences did not affect much the inference of tree topologies but yielded better branch length estimation when being larger, and the estimated median LUCA ages were closer to real values when the sampling time differences were not small despite larger variances, which was probably caused by a wider range of the simulated LUCA ages (Supplementary Figure 6). The mutation rates (Figure 4B) were slightly underestimated with fewer mutations and small sampling time differences and more accurately estimated otherwise. In summary, CNETML inferred phylogenies well when there was sufficient information in the data, with larger mutation rates or sampling time differences leading to higher accuracy. We also ran CNETML on haplotype-specific CNPs of data simulated with *dt* = 5 years and 1000 sites to compare with the results when using total CNPs (Figure 4C), but similar to our previous results in Figure 3, we did not observe large differences.

#### Performance on relative copy numbers

CNAs called from sWGS data with common tools, such as QDNAseq [21], are often relative values, which are hard to interpret, but they provide a way to mitigate the effect of WGD in phylogeny inference. For example, PISCA used a baseline strategy to convert absolute haplotype-specific copy numbers to relative values, which lead to better phylogeny inference and more accurate rate estimation on simulated data with WGD [3]. The basic idea is to divide the observed copy numbers by an estimated baseline (rounded mean copy number) for each haplotype and then round the values up or down randomly to reduce bias when the remainder is not zero. This is a simple strategy to process the absolute CNPs for reasonable phylogeny inference when WGD is present, as it is just one event changing ploidy, and the normalization by baseline copy number may cancel its effect. We adopted a similar strategy in CNETS to simulate relative haplotype-specific copy numbers by using the known ploidy as the baseline and relative total copy numbers by using 2^*N*^*WGD* as the baseline, where *N*_*WGD*_ is the number of WGD events in a genome. CNETS output simulated relative total CNPs by reducing the normalized copy numbers with the ploidy of the genome, with values smaller than −2 and larger than 2 set to −2 and 2 respectively for consistency with QDNAseq output.

We ran CNETML on relative total and haplotype-specific CNPs simulated with *c_max_* = 8, *dt* = 1 year, *u* = *e* = 0.001 per haplotype per site per year, and WGD rate 0.05 per year, which generated data of four types according to the distribution of WGD among samples: clonal WGD where WGD appears in all samples, multiple WGDs where there are more than one WGD across the tree but each sample has at most one WGD, single WGD where there is only one WGD across the tree, and no WGD. When running CNETML on relative total CNPs, we added the copy numbers with normal ploidy so that all values are positive. As a comparison, we also ran MEDICC2 [9], the only method to infer CNA-based phylogenies from NGS data at the presence of WGDs, on allele-specific CNPs which were converted from haplotype-specific CNPs by custom R scripts and CNETML on total CNPs respectively.

The results are grouped into four types by WGD distribution in the data (Figure 5). As expected, CNETML on absolute total CNPs reconstructed inaccurate phylogenies and missestimated mutation rates in most cases whenever WGD was present, with duplication rates largely overestimated especially on data with clonal WGD and deletion rates slightly underestimated. On data with single or multiple subclonal WGD(s), CNETML on relative CNPs can achieve similar performance in phylogeny inference to MEDICC2 on absolute allele-specific CNPs and the mutation rates were also accurately estimated with slight underestimation of deletion rates on relative haplotype-specific CNPs. On data without WGD, CNETML performed similarly on all types of data, which suggests using relative copy numbers still conserves the information for phylogeny inference and rate estimation. On data with clonal WGD, CNETML on relative CNPs reconstructed phylogenies less similar to the truth and underestimated mutation rates, particularly deletion rates on relative haplotype-specific CNPs, which is probably due to greater signal loss when converting copy numbers relative to a doubled ploidy. In summary, it seems entirely feasible to recover the phylogeny directly from relative copy numbers, such as those from QDNAseq, when WGD is not clonal. For further validation of the inference, empirical information or methods to detect WGD [54] or call absolute copy numbers [27] from sWGS data may be used to estimate the presence of clonal WGD.

**Figure 5:**
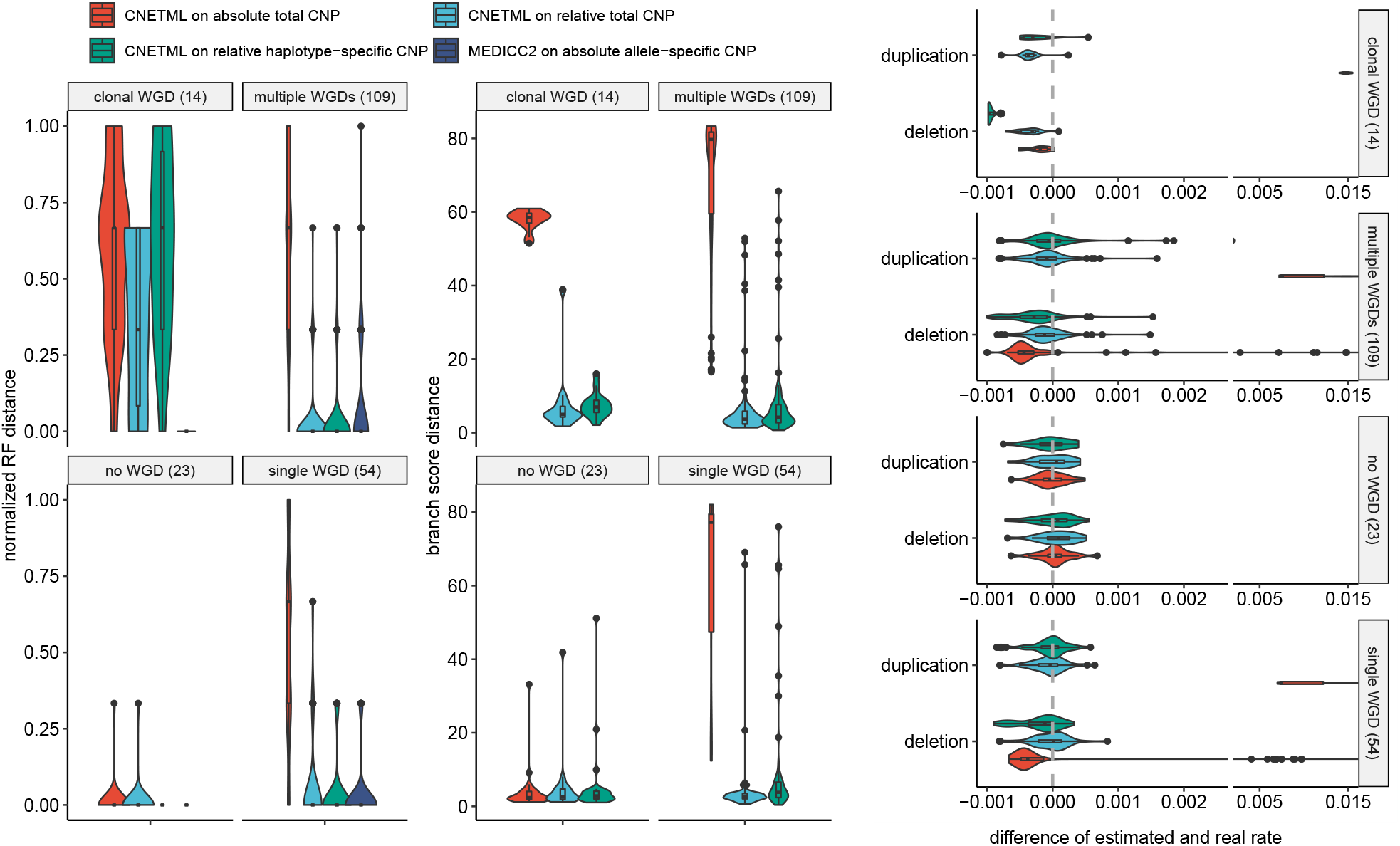
The performance of CNETML on relative copy number data. There are 200 simulated datasets in total, which are divided into four groups by the types of WGDs. The number of datasets in each group is shown in brackets. MEDICC2 was excluded when comparing branch score distance because the branch length in a tree built by it has a different meaning (the number of events between CNPs of two nodes based on CNT model) and it is hard to compare fairly. 29 outlier data points with values larger than the maximum of x-axis on datasets with subclonal WGDs are excluded in the plot of mutation rates for better visualization of the majority data. Box plots as those in Figure 3.

#### Performance under violation of independent sites assumption

ML inference of phylogenetic trees using sequence data was shown to be highly robust to violations of assumptions [37]. As our model of CNA evolution strongly depends on the independent sites assumption, we ran CNETS using the waiting time approach to generate duplications and deletions of different sizes to examine how overlapping CNAs affect the performance of CNETML. We simulated trees with *dt* = 1 year so that rate estimation is feasible and introduced duplications/deletions along the tree with rate *u* = *e* = 0.001 per haplotype per site per year and mean size being 1, 10, and 100 bins (500 Kbp, 5 Mbp, and 50 Mbp), respectively. These sizes were chosen because focal CNAs are typically defined as CNAs of size no larger than 3 Mbp [55] and 50 Mbp is larger than p-arm size of 15 autosomes and q-arm size of 4 autosomes to include arm-level CNAs. We built trees with CNETML using original bin-level data (site as bin) and post-processed segment-level data (site as segment, see section 4 for details of the post-processing).

As expected, the inferred phylogenies and mutation rates were more dissimilar to the ground truth with larger CNA sizes due to information loss as a result of overlaps, with overestimated branch lengths and mutation rates and slightly underestimated LUCA age (Figure 6A). However, when the mean duplication/deletion size was 5 Mbp, shorter than the typical focal CNA size, the bias was not very large, and the tree topologies were still recovered well. In addition, when we scaled the estimated rates by the mean duplication/deletion size, the estimation errors were much smaller (Figure 6B). These results suggest that slight violation of the independent sites assumption seems acceptable in phylogeny inference, and the estimated mutation rates may be scaled to account for the size of CNAs. On the other hand, the differences of using bin-level and segment-level data were small because the site patterns in the input data were similar in both cases, except that bin-level data might contain more sites and a larger number of the same pattern which lead to longer computing time. The rates and LUCA age were slightly overestimated with bin-level data on larger CNAs, which is probably due to larger overdispersion caused by dependencies between adjacent bins with the same copy numbers [56].

**Figure 6:**
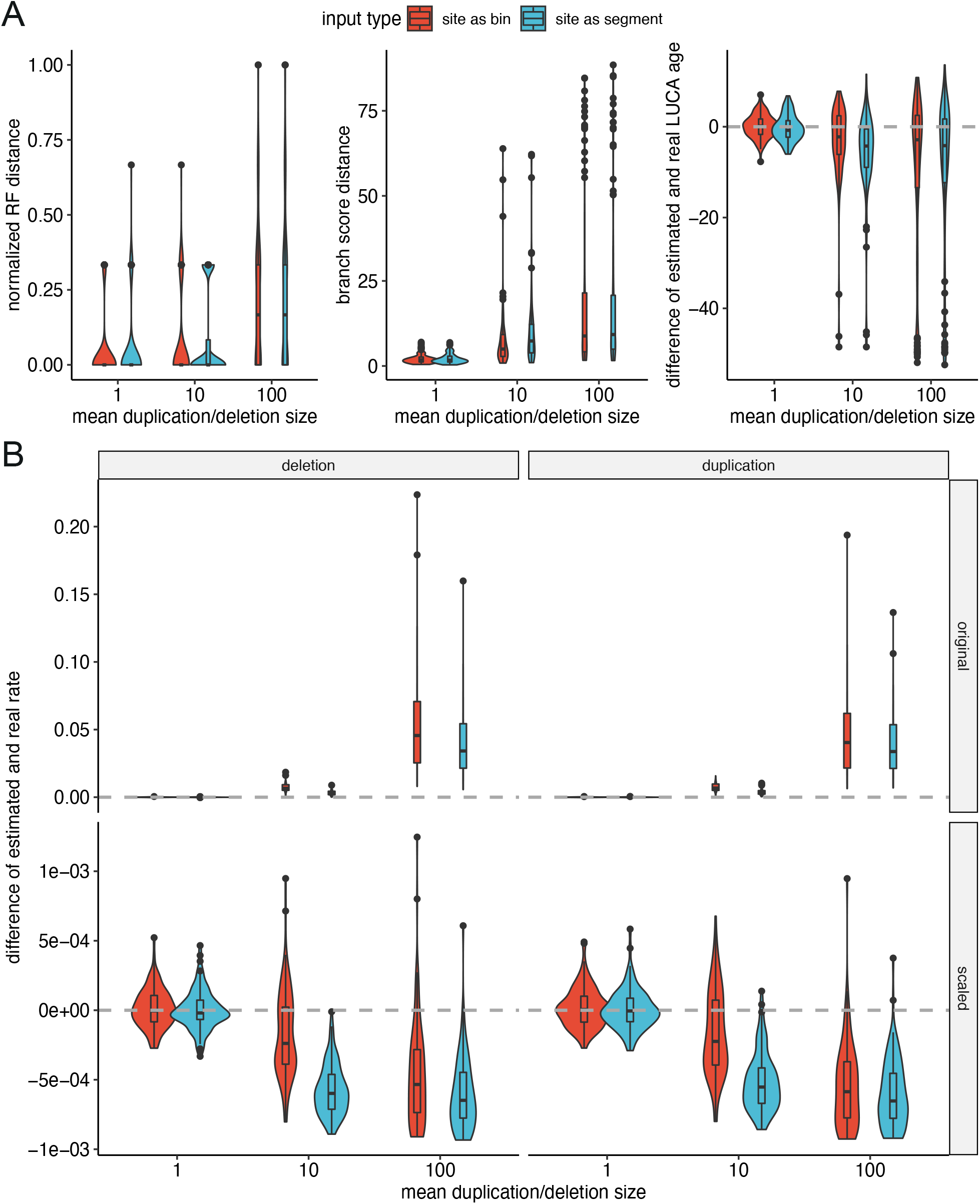
The performance of CNETML with different types of sites on data simulated with mean duplication/deletion of different sizes. **A**: The accuracy of phylogeny inference. **B**: The accuracy of mutation rate estimation before and after scaling. There are five samples in each simulated tree and 100 datasets for each parameter setting. Box plots as those in Figure 3.

Therefore, we recommend using segment-level data for faster computation, less correlation between segments, and better interpretability in practice.

### Application to Barrett’s esophagus patients

To demonstrate the applicability of CNETML on real data, we applied it to data for two BE patients in Figure 1 of [13], where CIN was used to predict risk progression (Figure 7). QDNAseq was applied on sWGS data to get relative CNPs in 589 bins of fixed size (about 5 Mbp) for each patient, which were normalized across the cohort of 777 endoscopy samples from 88 patients. One nonprogressor patient, 51, has 15 samples taken from 2006 to 2011, which shows similar CNPs across samples. The other progressor patient, 20, has 12 samples taken from 1998 to 2008, which shows more copy number variation across samples. Although WGD was shown to be prevalent in BE patients [3], it seems less likely to have clonal WGDs for these two patients given the large span of sampling times and diverse sampling locations. We rounded the provided fractional copy numbers to the nearest integers, set those smaller than −2 to −2 and larger than 2 to 2, and merged consecutive bins with the same copy numbers across all samples into segments. Since the exact patient ages were not provided, the patient age at the first sampling time was set to be 60 for patient 51 and 62 for patient 20, the mean age of all nonprogressors and progressors in the cohort at diagnosis of BE, respectively, which provides good approximations of the upper bounds of the tree heights during optimization.

**Figure 7:**
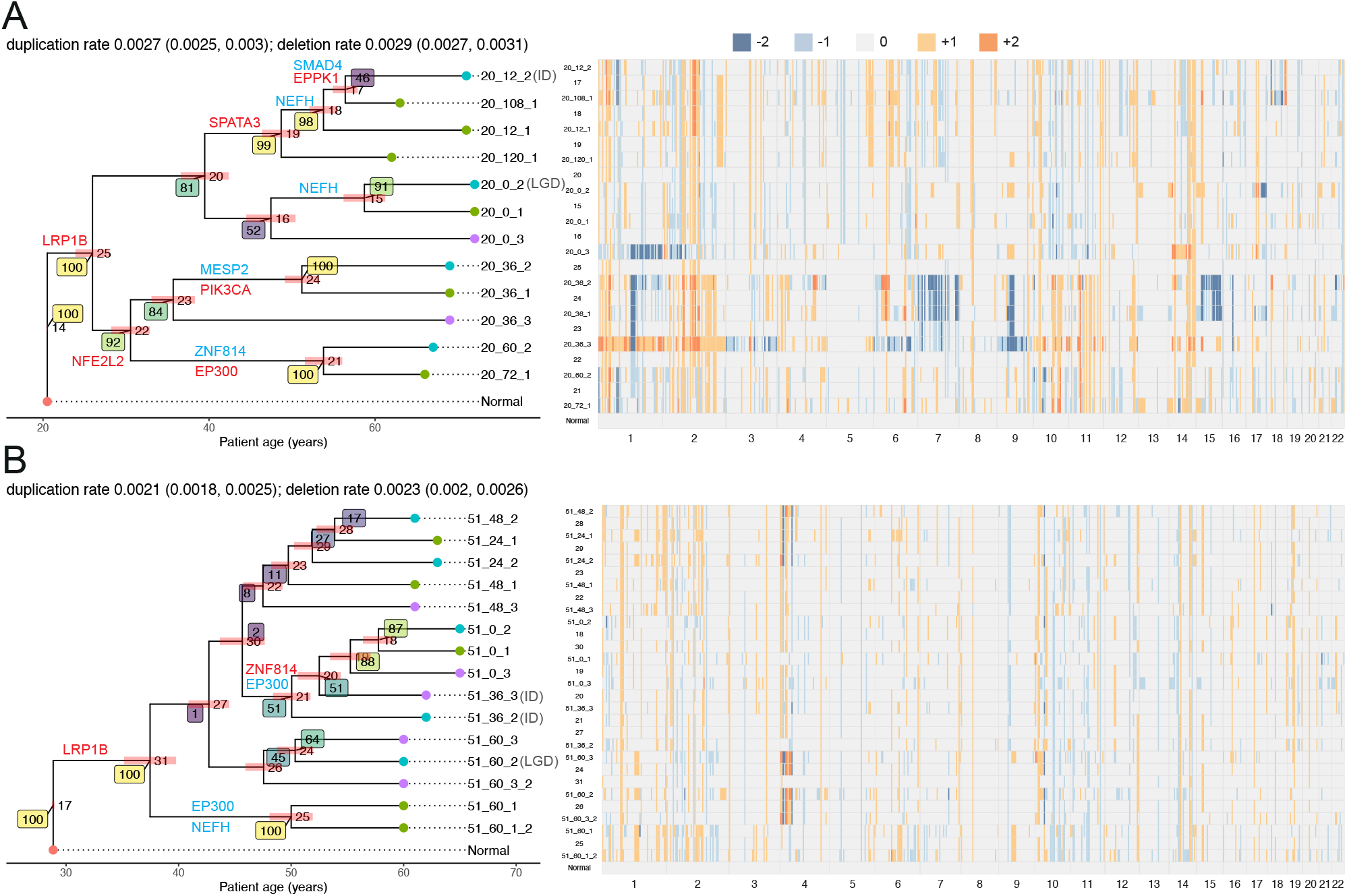
The ML trees and ancestral states reconstructed by CNETML for patient 20 (**A**) and 51 (**B**). The bootstrap support values are shown in coloured rectangles with lighter colours suggesting stronger support. The coloured bars at the internal nodes show the confidence intervals of node ages. Each sample is denoted by “patientID_timeID_locationID”, where timeID is the month before final endoscopy and locationID is the relative esophageal sample location. There are two sets of samples taken at the same time and location for patient 51, which are indicated by “_2” at the end. For patient 20, the samples taken at 12 months before final endoscopy location 2 (20_12_2) is Indeterminate (ID), the final endoscopy at location 2 (20_0_2) is Low-Grade Dysplasia (LGD), and all the other samples are Non-dysplastic BE (NDBE). For patient 51, the samples taken at 36 months before final endoscopy location 2 and 3 (51_36_2 and 51_36_3) are IDs, the sample taken at 60 months before final endoscopy at location 2 (51_60_2) is LGD, and all the other samples are NDBE. The cancer-related genes overlapping with the reconstructed CNPs are shown on the branches (red: copy number gain, blue: copy number loss). The confidence intervals of duplication/deletion rates are shown in parentheses in the title of the plot for each patient.

We first ran CNETML 100 times on the input data, selected the tree with largest likelihood, *T_b_*, and did 100 bootstraps to get branch support values for *T_b_*. Then we fixed the tree topology to *T_b_* and optimized node ages and mutation rates to get 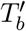, with the initial mutation rates set to the estimated rates on *T_b_*. We ran another 100 nonparametric bootstraps with the topology of 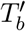 to get the confidence intervals (2.5th and 97.5th percentile) of node ages and mutation rates in 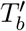. Lastly, we reconstructed ancestral CNPs based on 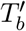 and checked the biological significance by computing their overlap with cancer-related genes from COSMIC Cancer Mutation Census (CMC) with keyword “oesophag” in the description of disease [57] and 75 regions selected by the elastic-net regression model as being predictive of BE progression (predictive regions) in [13].

The tree topology for patient 20 had bootstrap support values of more than 80% except for two branches (Figure 7A). Although the branch connecting the samples taken at 12 months location 2 and 108 months location 1 (times before final endoscopy) had only 46% of support, they shared a loss of gene SMAD4, which was shown to promote tumorigenesis from BE toward esophageal cancer [58] and hence suggested the reliability of this branch. The tree topology for patient 51 had much poorer support due to the lack of changes in copy numbers (Figure 7B). The estimated mutation rate of patient 20 was slightly higher than that of patient 51, around 0.006 and 0.004 per haplotype per site per year respectively, which is as excepted because progressors tend to have higher mutation rates and seems consistent with previous results for BE patients [3]. The LUCA age approximates the onset time of BE, since CNAs are likely to begin after BE establishes in the esophagus. The results suggest patient 20 had BE about 40 years before the first sample, about 10 years earlier than patient 51, which also seems consistent with previous results where two progressors had younger LUCA ages than two nonprogressors [3]. The estimated phylogenies also show a longer dwell time (the time a patient has lived with the precursor) of BE in the progressor (patient 20) than the non-progressor (patient 51), which is consistent with previous modeling results [59]. From the reconstructed CNPs of the MRCAs of both patients (node 25 for patient 20 in Figure 7A and node 31 for patient 51 in Figure 7B), we found gene LRP1B included in a region on chr 2q with copy number gain (see Supplementary Table 2 for the complete list of overlaps). The original average relative copy number for patient 20 (1.4) across all samples in this region is about twice that for patient 51 (0.6), suggesting more gains in patient 20. Although most common alterations involving LRP1B are simple somatic mutations or copy number losses, 4.89% cases have copy number gains in the TCGA-ESCA cohort with 185 cases [60]. The CNP of the MRCA of patient 20 also had a region of gain on chr 4, which overlapped with the predictive region whose associated coefficient of variation for the relative risk (CV) is 1.018 (ranked 15th among 75 regions) [13]. The CNPs of the MRCA of patient 51 and the ancestors of the top and bottom lineages of patient 20 (node 20 and 22 in Figure 7A) overlapped with the predictive region whose associated CV is 5.090 (ranked 6th among 75 regions) [13]. The CNP of the ancestor of the bottom lineage of patient 20 also had a region of gain overlapping with gene NFE2L2 on chr 2q, which has about 11.41% cases with copy number gains in TCGA-ESCA cohort [60]. For patient 51, the lineage starting from node 21 had a region of gain overlapping with gene ZNF814, which has 13.04% cases with copy number gains in TCGA-ESCA cohort [60]. All these findings suggest that the phylogenies and mutation rates inferred by CNETML are biologically meaningful and additional insights into carcinogenesis can be gained from the reconstructed ancestral CNPs.

## 3 Discussion

In summary, we developed CNETML, a new ML method to reconstruct the evolutionary history of multiple samples taken from a single patient at different locations and/or times, which can take as input (relative) total integer copy numbers called from sWGS data. CNETML is capable of jointly estimating the node ages and mutation rates by year when patient samples were taken at different times. The estimation provides approximate timing of initiating CNAs and hence possible onset age of the disease such as BE in a patient, which cannot be detected clinically due to asymptomaticity but is important in carcinogenesis and may be helpful in cancer screening and surveillance programs [59]. This capability is derived from a novel Markov model of CNA evolution, which assumes the sites (bins or segments) in a CNP are independent and hence allows the usage of classical methods for phylogeny inference and ancestral reconstruction. We evaluated CNETML on data simulated with CNETS, our novel program of general utility to simulate CNAs along a phylogenetic tree. The simulations suggest that CNETML performed well when there were sufficient CNAs and/or timing information in the data, even on relative CNPs with subclonal WGDs. The ability to work on relative CNPs makes CNETML applicable to a wide range of sWGS data obtained from cancer patients, which was demonstrated by its application on two BE patients, where we inferred sample phylogenies along with ancestral CNPs which suggest the time LUCA arose and early CNAs driving the malignancy. Although caution is still required when interpreting the inference on relative CNPs without knowing the exact presence of clonal WGDs, the inference on relative copy numbers provide a reference for further improvement. CNETML is also applicable to allele-specific CNPs if they have been phased to distinguish haplotypes, and the performances were similar to those on total CNPs when there were less than 10% of copy-neutral CNA events (such as cn-LOH and MSAI) across all sites. Despite the independent sites assumption, CNETML was robust to considerable amounts of overlaps among simulated focal CNAs.

Although CNETML aims to build a sample tree where each tip is a CNP from a patient sample, it can be used to build trees from CNPs of subclones or single cells, since the main input is simply an integer copy number matrix where the rows can represent subclones or cells. The input CNPs are assumed to be called from sWGS data and hence cover the whole genome, but it may be applicable to SNP array or WES data if the gaps between segments with atypical copy numbers are filled to avoid acquisition bias [61].

In principle, the likelihood-based approach adopted in CNETML is more sophisticated than distance matrix and MP methods. To deal with the specific properties of (relative) CNPs and allow for more flexible evolutionary models specific for carcinogenesis, we implemented a novel tool rather than using existing frameworks designed for traditional phylogenetic inferences, such as BEAST [40, 41]. Future development of the model could include Markov chains at different scales to incorporate chromosomal and/or arm level gain/loss and WGD, and the use of penalized maximum likelihood estimation to incorporate prior knowledge on parameters when there are insufficient information in the data [62]. Another development would be the estimation of varying mutation rates in different lineages under a relaxed local clock [63]. Finally we can extend to a fully Bayesian approach, which can impose informative prior distributions and naturally provide a measure of uncertainty of the inference (posterior probabilities of sampled trees) despite a higher computational cost.

The inference of sample phylogeny from CNAs called from sWGS data is a very challenging problem. Although CNETML makes progresses in tackling some issues, it still has a few limitations in data handling. First, we assumed the input CNPs are complete and accurate, which is often violated in reality. Errors in copy number calls, which may arise from poor calling or missing data, directly affect the inference, since just one wrong copy number called at a site of a sample may lead to a unique site pattern and bias likelihood computation. Methods developed for sc-seq data often incorporate approaches to deal with data noise [19], which may be extended to CNAs called from sWGS data, such as combining CNA calling from raw read counts with phylogeny inference [5, 7] and incorporating false positives and false negatives into the model directly [6]. Moreover, we assume each sample is homogeneous with only one clone and do not deal with clonal deconvolution. This is reasonable to some extent as CNAs detected from sWGS data typically represent the dominant clone in a sample, which is different from sample trees built from SNVs that often represent highly admixed cell lineages [64]. However, given data of higher resolution, it would be helpful to quantify ITH and how it affects the monoclonal assumption.

In summary, we have provided a tool that can enhance the use of sWGS and allow for spatio-temporal inferences of carcinogenesis in patients. Due to the relatively low cost of sWGS, we believe our approach will have increasing impact in understanding the biology of carcinogenesis and will underlie future clinical applications.

## 4 Methods

### Preprocessing of input data

The input CNPs for CNETML are mainly obtained from common CNA calling methods for sWGS data. For example, QDNAseq [21] is often used to get relative copy numbers by computing read counts in fixed-sized bins, doing segmentation, and calling copy numbers with CGHcall [65] which classifies copy numbers into: double deletion (−2), single deletion (−1), normal (0), gain (1), and amplification (2). To get the data matrix *D*, we assumed the same binning or segmentation across all samples to get consistent sites. When raw copy number calls were at bin level, segments were obtained by merging consecutive variant bins on the same chromosome with the same copy number across all samples.

The input sampling dates were converted to years (divided by 365). For convenience, the time for the first sample was set to 0 and the time for other samples was then counted as the number of years starting from the first sample.

### The computation of likelihood

The probabilities of observing data at a single site *P* (*D*^(*i*)^|*T, θ*) was computed with Felsenstein’s pruning algorithm by post-order traversal of *T*, in which each node is visited only after all its descendants have been visited [43]. The computation can be expressed as a recursion that computes *L_i_*(*d_i_*) for each node *i* at each possible haplotype-specific copy number state *d_i_*, the conditional probability of observing data at the descendant tips below *i*. Let *p_d_idj* (*t_j_*) represents the transition probability of *d_i_* becoming *d_j_* after time *t_j_*, where *i* and *j* are two nodes in *T* connected by a branch of length *t_j_*. Suppose the root (LUCA node) is *r* with state *d_r_* = 4 since it is assumed to have normal diploid copy number, then:

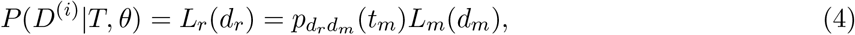

where *m* is the MRCA node connected to *r* with a branch of length *t_m_*. When node *i* is an internal node with children node *j* and *k*,

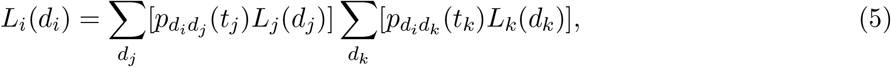

where *t_j_* and *t_k_* are the lengths of the branch from node *i* to *j* and *k* respectively. When node *i* is a tip with observed haplotype-specific copy number state *c_i_*,

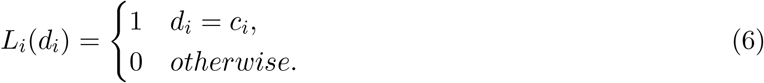

When the input copy numbers are total, there may be multiple haplotype-specific copy number states compatible with the observed value at tip *i*, denoted by set *S_i_*, and hence

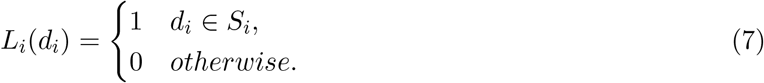

To improve efficiency in likelihood computation, the transition probability matrix, *P* (*t_j_*) = *e^Qt^j*, was computed once with scaling and squaring method [66] for each branch of length *t_j_* and used for all sites. *L_r_*(*d_r_*) for two sites with the same site patterns were computed once too, as they have the same probability of being observed.

### Statistical phylogeny inference with maximum likelihood method

An important aspect in optimization of likelihood function *L*(*T*) is the incorporation of bound constraints in L-BFGS-B algorithm. To avoid negative branch length, we define a minimal branch length *l_m_* (1e-3 year by default). To encode the constraints imposed by patient ages at different sampling times, we define a new variable *x_i_* for an internal node *i* with child *j* on a tree *T* of *n* samples:

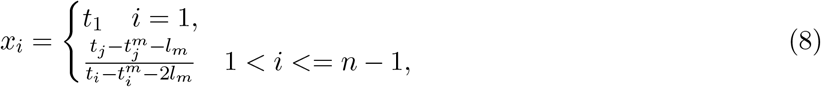

where *i* is from 1 (the root) to *n* − 1, *t_i_* is the age of node *i*, and 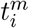 is the maximum age of the tips below node *i*. Because the parent age should always be smaller than those of the children nodes, *x_i_* has bounds as below:

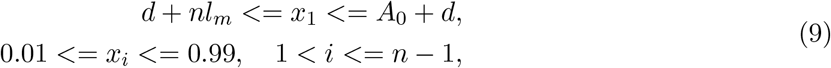

where *A*_0_ is the patient age at the first sample and *d* is the time difference between the last and first sample.

For exhaustive tree search, we enumerated all the possible tree topologies for the given number of samples and then found the ML tree by optimizing the parameters. For heuristic tree search, we started with a number of initial trees (100 by default), selected those with unique topologies, and computed their approximate likelihoods. Then we selected the top *n*_1_ (20 by default) trees ordered by decreasing likelihoods to do hill-climbing nearest neighbor interchanges (NNIs) [45] and kept the top *n*_2_ (5 by default) trees with largest likelihoods for further optimization to get the ML tree. To avoid local optima, we built parsimony-based stepwise addition trees as initial trees, which were obtained by using function random.addition in R library ape [48] and transformed into the formats acceptable by CNETML.

### Data simulation

The overall procedure of simulations in CNETS is as follows:

1. Generate a random coalescence tree of *n* samples. Available trees can also be given as input.
2. Optionally simulate temporal samples for a patient of specified age. One way to generate samples at various time points with just one additional parameter, *dt*, is as below. Note that this simulation approach destroys the coalescence structure but it is sufficient to generate a sample phylogeny for the purpose of testing phylogeny inference methods like CNETML.

a. Assign random times (in year) to the tips by changing terminal branch lengths with multiples of *dt*.
b. Rescale the internal branches of the tree so that the tree height is no larger than the patient age at the last sampling time.
3. Simulate CNPs on the tree with the Markov model of CNAs.

a. Generate the CNP for the root (normal diploid genome).
b. Simulate CNPs directly at the end of each branch according to the transition probability matrix or simulate mutational events along each branch by using exponential waiting times.
4. Output result files.

When simulating CNPs directly given the total number of sites (segments), we distributed the sites roughly according to the size of each chromosome with Dirichlet distribution. Each genome with *m* sites was represented by its CNP (*c*_1_, *c*_2_, …, *c_m_*) whose initial values at all sites are 2 for total copy number data or 4 for haplotype-specific copy number data. For a site *i* with state *c_i_*, we sampled its target state from the discrete distribution specified by row *i* of transition probability matrix *P* (*l*) for a branch of length *l*.

When simulating events of multiple scales by waiting times, we pre-specified the number of sites (bins) on each autosome of the reference genome with an array [367, 385, 335, 316, 299, 277, 251, 243, 184, 210, 215, 213, 166, 150, 134, 118, 121, 127, 79, 106, 51, 54], which were extracted from QDNAseq output on real data with 4,401 bins of 500 Kbp. Each genome with *m* sites was initially represented by the set of sites, denoted by *G* = (*l*_1_, *l*_2_, …, *l_m_*). We denoted the diploid genome by *G_d_* = [*G, G*], which was implemented by making a copy of *G* to represent the other haplotype. The final CNP of the genome (*c*_1_, *c*_2_, …, *c_m_*) was computed for each site by adding up the number of copies across all the haplotypes when considering total copy number or across the specific haplotype when considering haplotype-specific copy number. Some constraints were imposed to get more realistic data: 1) Chromosomal gain and WGD were only possible when the resultant maximum copy number is smaller than the specified *c_max_*; 2) The duplication/deletion stopped at the end of a chromosome. For the simulation of specific mutational events along a branch of length *l* from initial time *t* = 0, we used the following steps:

1. Generate a random waiting time *e* from the exponential distribution with rate *r*, where *r* is the total mutation rate across the genome, obtained by adding up the duplication and deletion rates across all sites along the genome, chromosomal gain and loss rates across all chromosomes, and WGD rate.
2. Generate a mutation, whose type is randomly chosen based on the relative rates of different event types.

a. For segment duplication/deletion, randomly choose the start bin based on the rates across sites, the haplotype, and the size in the number of bins, where a duplication can be either tandem (duplicated at the end of the current location) or interspersed (inserted in any position across the genome) with equal possibilities.
b. For chromosome gain/loss, randomly select the chromosome according to the rates across chromosomes and the haplotype.
3. *t* = *t* + *e*.
4. Stop when *t* >= *l*.

## 5 Acknowledgements

CPB and BL acknowledge funding from the Wellcome Trust (209409/Z/17/Z). The authors acknowledge the use of the UCL Myriad High Throughput Computing Facility (Myriad@UCL), and associated support services, in the completion of this work. We thank Simone De Angelis, Rachel Muir, and Christos Magkos for testing our programs. We thank William Cross for helpful suggestions on the manuscript.

