## Supplemental Tables and Figures for "CNETML: Maximum likelihood inference of phylogeny from copy number profiles of spatio-temporal samples"

Supplementary Table 1: The rate matrix $Q$ when the maximum total copy number $c_{max} = 4$.

| | **0** | **1** | **2** | **3** | **4** | **5** | **6** | **7** | **8** | **9** | **10** | **11** | **12** | **13** | **14** |
|---|---|---|---|---|---|---|---|---|---|---|---|---|---|---|---|
| | **0/0** | **0/1** | **1/0** | **0/2** | **1/1** | **2/0** | **0/3** | **1/2** | **2/1** | **3/0** | **0/4** | **1/3** | **2/2** | **3/1** | **4/0** |
| **0/0** | 0 | 0 | 0 | 0 | 0 | 0 | 0 | 0 | 0 | 0 | 0 | 0 | 0 | 0 | 0 |
| **0/1** | $e$ | $-(u+e)$ | 0 | $u$ | 0 | 0 | 0 | 0 | 0 | 0 | 0 | 0 | 0 | 0 | 0 |
| **1/0** | $e$ | 0 | $-(u+e)$ | 0 | 0 | $u$ | 0 | 0 | 0 | 0 | 0 | 0 | 0 | 0 | 0 |
| **0/2** | 0 | $e$ | 0 | $-(u+e)$ | 0 | 0 | $u$ | 0 | 0 | 0 | 0 | 0 | 0 | 0 | 0 |
| **1/1** | 0 | $e$ | $e$ | 0 | $-2(u+e)$ | 0 | 0 | $u$ | $u$ | 0 | 0 | 0 | 0 | 0 | 0 |
| **2/0** | 0 | 0 | $e$ | 0 | 0 | $-(u+e)$ | 0 | 0 | 0 | $u$ | 0 | 0 | 0 | 0 | 0 |
| **0/3** | 0 | 0 | 0 | $e$ | 0 | 0 | $-(u+e)$ | 0 | 0 | 0 | $u$ | 0 | 0 | 0 | 0 |
| **1/2** | 0 | 0 | 0 | $e$ | $e$ | 0 | 0 | $-2(u+e)$ | 0 | 0 | 0 | $u$ | $u$ | 0 | 0 |
| **2/1** | 0 | 0 | 0 | 0 | $e$ | $e$ | 0 | 0 | $-2(u+e)$ | 0 | 0 | 0 | $u$ | $u$ | 0 |
| **3/0** | 0 | 0 | 0 | 0 | 0 | $e$ | 0 | 0 | 0 | $-(u+e)$ | 0 | 0 | 0 | 0 | $u$ |
| **0/4** | 0 | 0 | 0 | 0 | 0 | 0 | $e$ | 0 | 0 | 0 | $-e$ | 0 | 0 | 0 | 0 |
| **1/3** | 0 | 0 | 0 | 0 | 0 | 0 | $e$ | $e$ | 0 | 0 | 0 | $-2e$ | 0 | 0 | 0 |
| **2/2** | 0 | 0 | 0 | 0 | 0 | 0 | 0 | $e$ | $e$ | 0 | 0 | 0 | $-2e$ | 0 | 0 |
| **3/1** | 0 | 0 | 0 | 0 | 0 | 0 | 0 | 0 | $e$ | $e$ | 0 | 0 | 0 | $-2e$ | 0 |
| **4/0** | 0 | 0 | 0 | 0 | 0 | 0 | 0 | 0 | 0 | $e$ | 0 | 0 | 0 | 0 | $-e$ |

Supplementary Table 2: Parameters used for tree generation in CNETS.

| | | |
|---|---|---|
| effective population size | $N_e$ | 90000 |
| generation time in year (365 days) | $t$ | 0.002739726 |
| exponential growth rate | $\beta$ | 1.563e-3 |

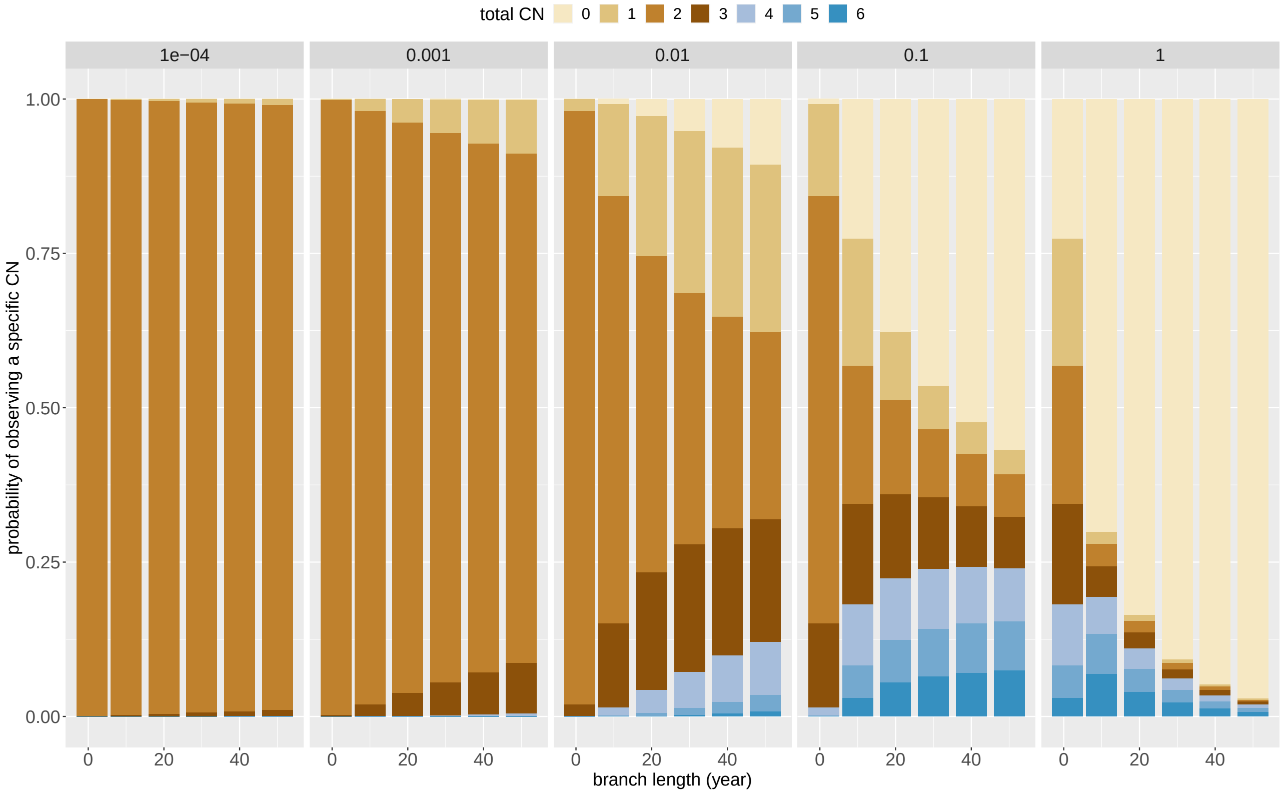

Supplementary Figure 1: The distribution of total copy number (CN) at the end of one branch under the Markov model. The plots are grouped by mutation rates. In each group, the x-axis shows branch length at size 1, 10, 20, ..., 50. The y-axis shows the probability of changing from normal total copy number (2) to each possible total copy number. We computed the final states of the Markov chain for one branch of varying lengths starting at normal state, copy number (1,1). When the mutation rate is very low (1e-4 per haplotype per site per year), there are only a few mutations and most sites stay normal. When the mutation rate is high (1 per haplotype per site per year), more sites reach absorbing states (copy number 0).

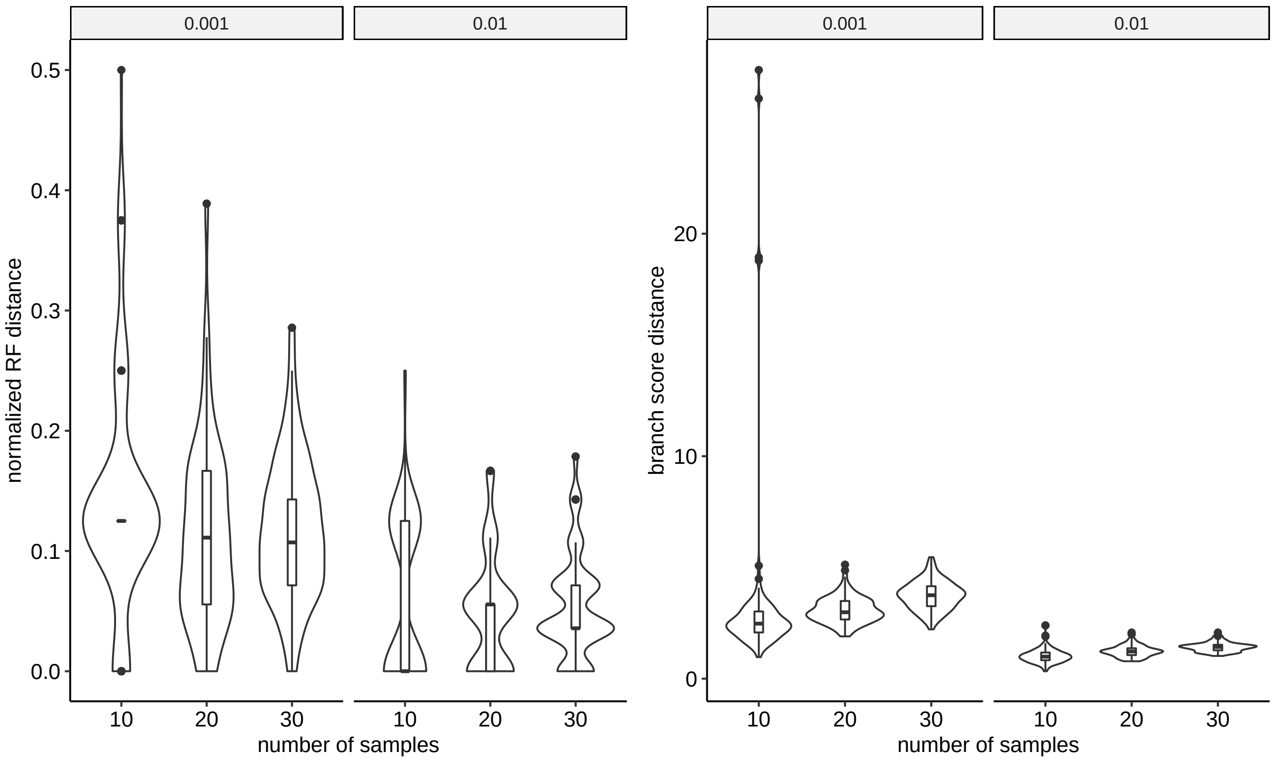

Supplementary Figure 2: The performance of CNETML (stochastic tree search) on data simulated with different mutation rates and number of samples. All the simulated samples are at the same time. The plots are grouped by mutation rates. There are 100 datasets for each parameter setting. The box plots show the median (centre), 1st (lower hinge), and 3rd (upper hinge) quartiles of the data; the whiskers extend to 1.5× of the interquartile range (distance between the 1st and 3rd quartiles); data beyond the interquartile range are plotted individually.

Supplementary Table 3: The data simulated by CNETS under different temporal signal strengths, grouped by the mean pairwise absolute difference of tip relative times.

| group | mutation rate (per haplotype per site per year) | number of sites | number of samples |
|---|---|---|---|
| small | 0.001 | 1000 | 103 |
| | | 10000 | 106 |
| | 0.01 | 1000 | 106 |
| | | 10000 | 103 |
| intermediate | 0.001 | 1000 | 108 |
| | | 10000 | 99 |
| | 0.01 | 1000 | 104 |
| | | 10000 | 108 |
| high | 0.001 | 1000 | 89 |
| | | 10000 | 95 |
| | 0.01 | 1000 | 90 |
| | | 10000 | 89 |

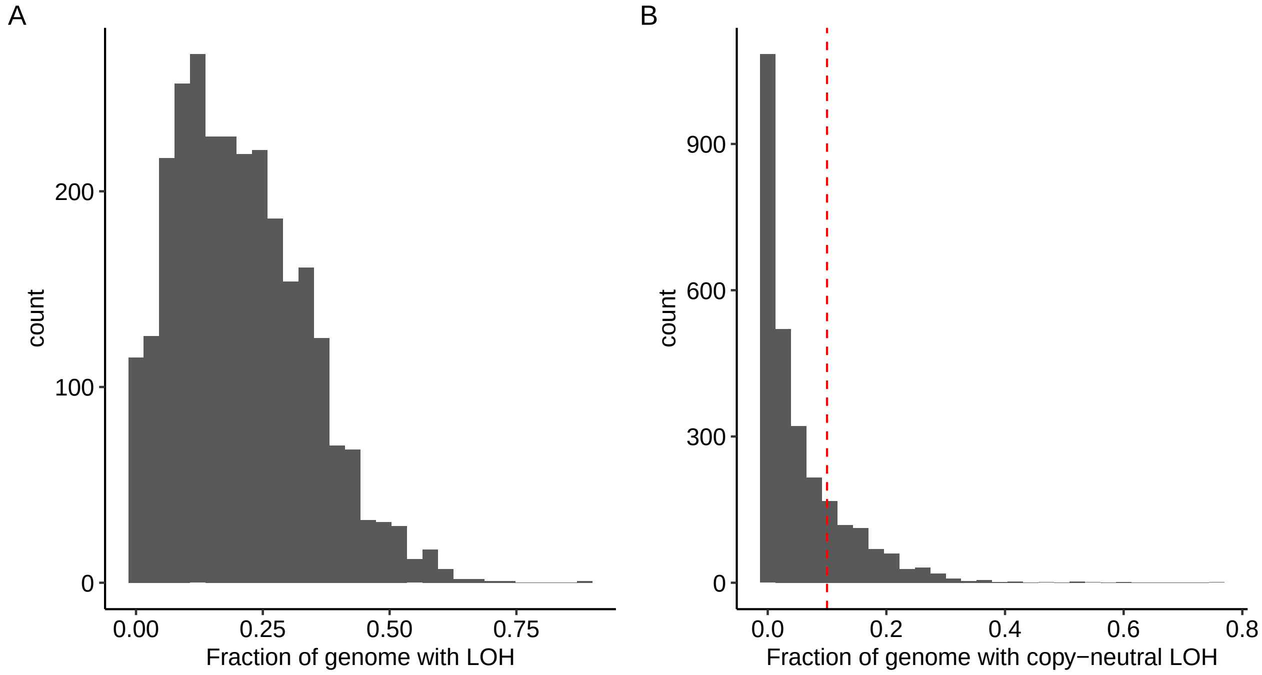

Supplementary Figure 3: The distribution of the fractions of genome with loss of heterozygosity (LOH) on 2778 samples from PCAWG dataset. **A**: The distribution of the fractions of genome with LOH. **B**: The distribution of the fractions of genome with copy-neutral LOH (red dashed line: 0.1).

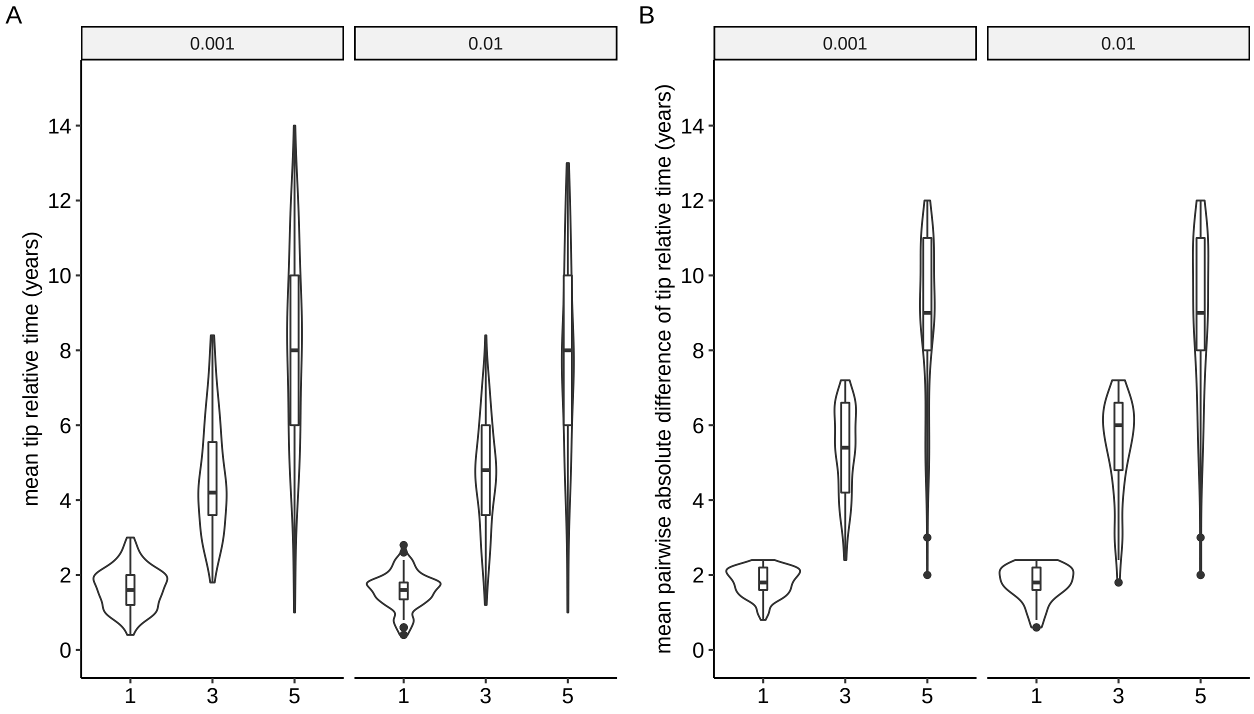

Supplementary Figure 4: The range of simulated sampling times under different temporal signal strengths and mutation rates. **A**: The average of the relative times at the tips (assuming the first sample is at time 0) in the simulated trees. **B**: The average of pairwise absolute difference of the relative times at the tips in the simulated trees. The x-axis shows the value of $dt$ which controls temporal signal strength, with larger value indicating larger time differences among samples. The plots are grouped by mutation rates. There are five samples in each simulated tree and 100 datasets for each parameter setting. Box plots as those in Supplementary Figure 2.

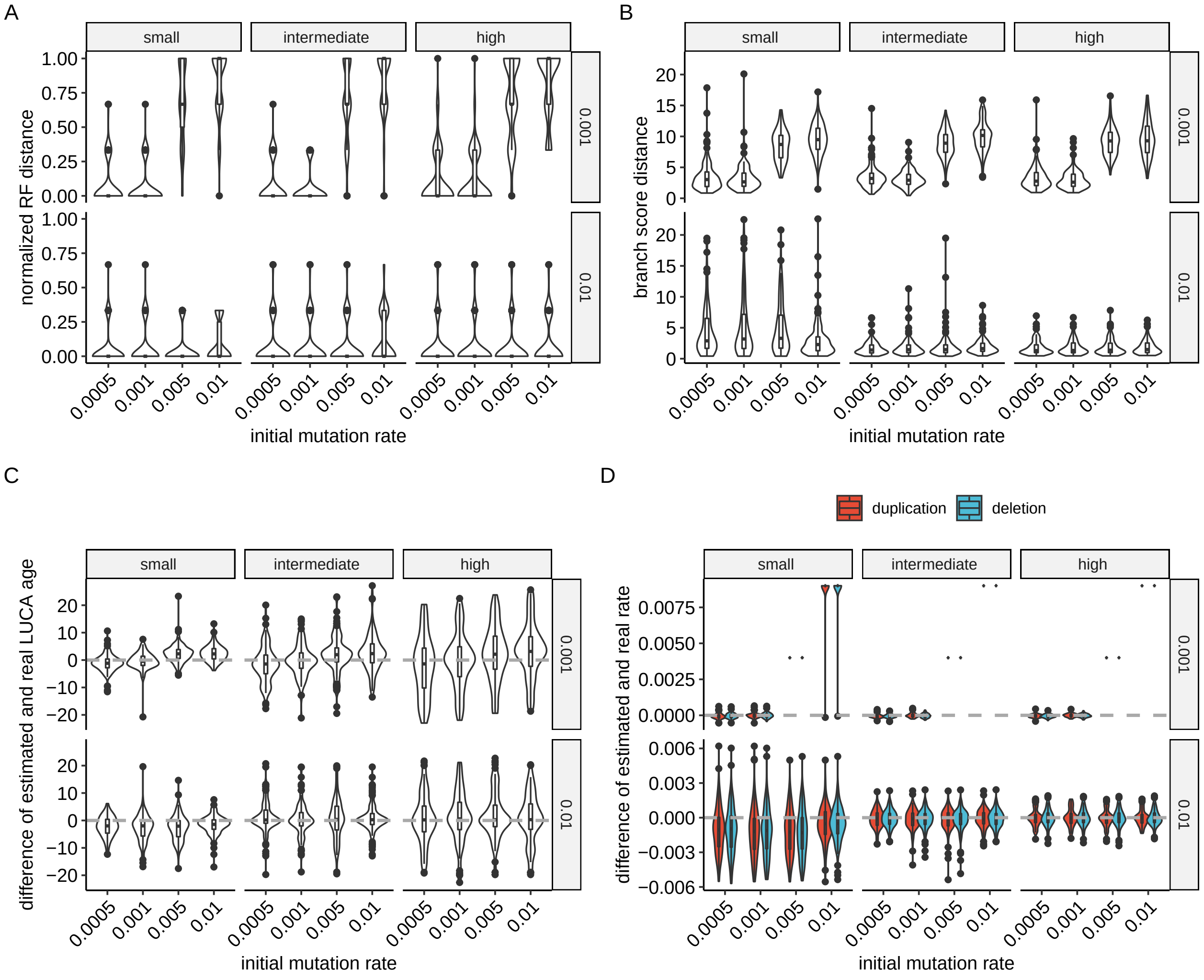

Supplementary Figure 5: The sensitivity of CNETML to initial mutation rates when jointly estimating the tree topology, node ages, and mutation rates on data simulated with different temporal signal strengths and mutation rates. **A-C**: The accuracy of tree inference under different initial mutation rates. **D**: The accuracy of mutation rate estimation under different initial mutation rates. There are five samples in each simulated tree and 100 datasets for each parameter setting. Box plots as those in Supplementary Figure 2.

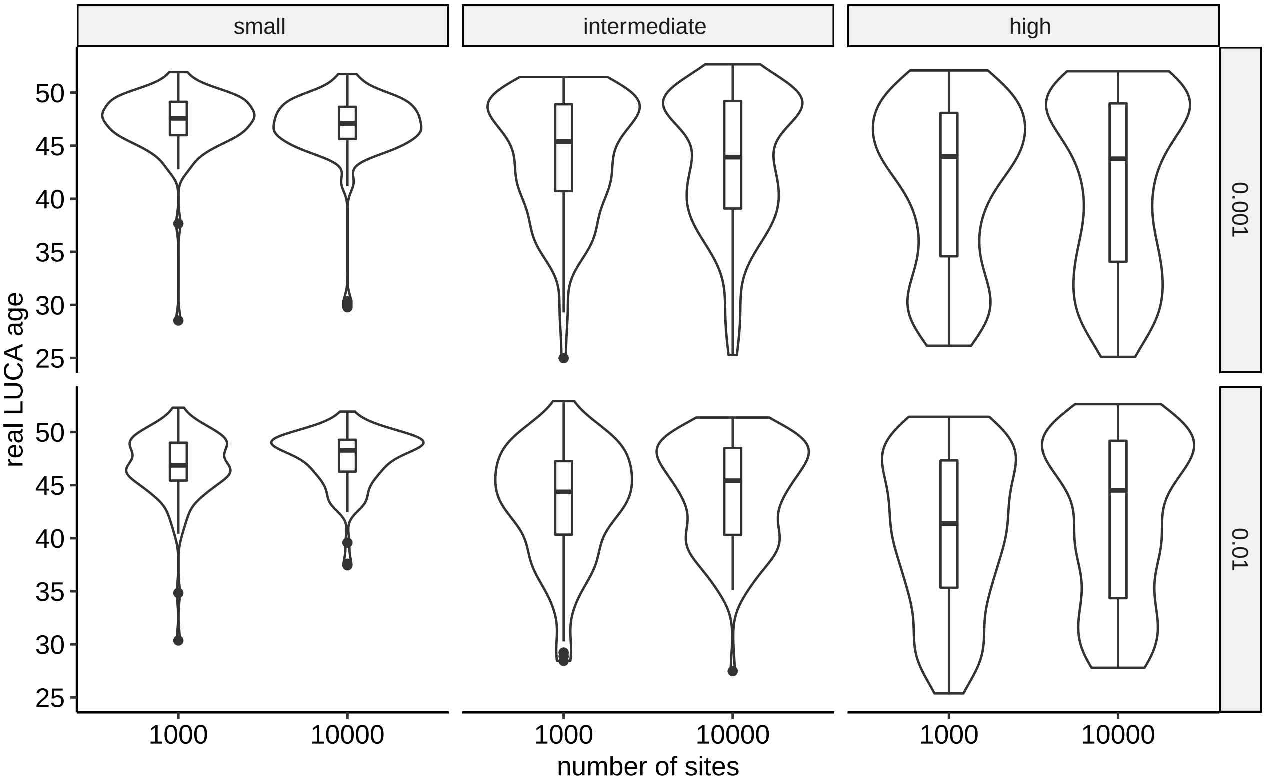

Supplementary Figure 6: The simulated real LUCA ages under different temporal signal strengths and mutation rates. The plots are grouped by mutation rates and sampling time differences. There are five samples in each simulated tree and 100 datasets for each parameter setting. Box plots as those in Supplementary Figure 2.
